# Model Selection for Biological Crystallography

**DOI:** 10.1101/448795

**Authors:** Nathan S. Babcock, Daniel A. Keedy, James S. Fraser, David A. Sivak

## Abstract

Structural biologists have fit increasingly complex model types to protein X-ray crystallographic data, motivated by higher-resolving crystals, greater computational power, and a growing appreciation for protein dynamics. Once fit, a more complex model will generally fit the experimental data better, but it also provides greater capacity to overfit to experimental noise. While refinement progress is normally monitored for a given model type with a fixed number of parameters, comparatively little attention has been paid to the selection among distinct model types where the number of parameters can vary. Using metrics derived in the statistical field of model comparison, we develop a framework for statistically rigorous inference of model complexity. From analysis of simulated data, we find that the resulting information criteria are less likely to prefer an erroneously complex model type and are less sensitive to noise, compared to the crystallographic cross-validation criterion *R*_free_. Moreover, these information criteria suggest caution in using complex model types and for inferring protein conformational heterogeneity from experimental scattering data.

## I. INTRODUCTION

Refinement of a Protein Data Bank (PDB) formatted model is intended to determine the molecular conformation(s) that best explain the diffraction patterns of X-rays from a macromolecular crystal. There is growing interest in fitting increasingly complex types of models to the experimental data, because some protein crystals can now be resolved at *Å* ngstrom scales and finer [1–3]; increasing automation and computational power have made it more tractable to parameterize more complex models; and finally, there is an emerging appreciation for the dynamic behavior of proteins *in vivo* [4–11].

At its simplest, a PDB structural model consists of a set of *x, y, z* coordinates for each atom. In addition, the B-factor (atomic displacement parameter) can be assigned at the level of each atom, residue, chain, or unit cell. Further complicating this is the anisotropic B-factor [12], which is defined by 6 parameters and can similarly be assigned per atom or at coarser groupings (*e.g.*, in the case of translation-libration-screw refinement [13]). More complex types of PDB structural models may include multiple *x, y, z* coordinates for an atom as alternative conformations with associated occupancies (“*q*”) [14]. Therefore, the parameters needed to describe a single atom in a crystal can vary from about three (when just *x, y, z* are refined) to ten times the number of conformations (when *x, y, z*, anisotropic B-factors, and occu-pancy are all refined per conformation). Although general best practices exist, quantifying when to increase the model complexity is currently an outstanding challenge.

A fundamental problem in statistics results from the fact that a more complex model with more parameters can typically be refined to fit data better than a simpler one [15]. This is necessarily true for nested model types, where particular constraints on the parameters of the complex model type reduce it to the simple model type. When the underlying physical reality is simple, the complex model will overfit the data—representing noise as physically meaningful structure [16]—despite reporting better quality of fit than the simpler model. Furthermore, the data may be limited (low resolution) and the potential space of complex solutions that equally satisfy the data may be large. Hence, practitioners need disciplined methods to decide when the data have both sufficient heterogeneity and sufficiently low noise to justify inferring more complex phenomena. We ask how much better a complex model type’s quality of fit should be to justify the additional parameters it requires beyond those of a relatively simple model type.

In crystallography, prevailing methods used to select among competing model types rely on the crystallographic reliability index or “*R*-factor” [17], an estimate of the error in the fitted crystallographic model:

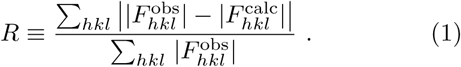

Here sums are taken over Miller indices *hkl* of the observed reflections’ amplitudes 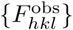 with respect to hypothetical ones 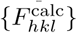 calculated from the crystal model [18].

The cross-validation statistic *R* _free_ is the *R*-factor calculated on randomly chosen test data, distinct from the working data used to fit the model parameters [19]. Statistical cross-validation is a general technique that judges a model type’s merit by quantifying its success at predicting test data that is excluded from use for refining the model type’s parameters, and thus reports on the predictive error [20, 21].

Nevertheless, cross-validation has limitations. For one, a cross-validation measure is by necessity calculated from a fraction of the total data set [19], reducing the set of measurements over which statistical fluctuations can be averaged (though this can be somewhat mitigated by *K-* fold cross validation [22] or its variant in the *R* complete method [23]). This increases relative noise, a particular concern for low-resolution refinement. Furthermore, cross-validation scores are often used to guide parameter refinement [24, 25] (*e.g.*, for global weights for geometry and B-factor restraints in Phenix [“Python-based Hierarchical ENvironment for Integrated Xtallography”] [26]). This practice presents some risk of producing models that have been overfit to the cross-validation measure, potentially diminishing its validating power [21]. This has led some to cross-validate the cross-validation measure itself using *R* sleep, calculated from a data fraction excluded from both refinement and cross-validation [27].

The risk of overfitting a model increases with the number *N* param of fitted parameters [28]. Model types (such as isotropic B-factors) with fewer parameters have less opportunity to overfit to noise than more complex model types (such as anisotropic B-factors). (For example, when fitting to *N* two-dimensional data points, an *N* th-order polynomial can fit perfectly but also captures any noise in the data, whereas a straight line is too simple to fit data with significant spread.) Yet fewer parameters give a model type less chance to capture real heterogeneity present in the data. The limitations of cross-validation raise basic practical questions.

How large a change in *R* _free_ is necessary to confidently adopt a more complex model type? Significance tests for *R*-factors have been proposed [29, 30] and employed to judge the suitableness of B-factor anisotropy [31], but don’t appear to be widely used. In statistics, such practical questions have generated a whole field of research on *model selection* [15, 16], which is intended to determine how to weigh the benefit of a more complex model type’s improved fit to data against its risk of overfitting, and to discern the appropriate level of model complexity to use to fit a particular data set.

Two simple and widely used model selection criteria are the Akaike Information Criterion (AIC) [28],

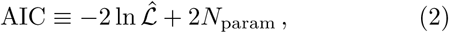

and the Bayesian Information Criterion (BIC) [32],

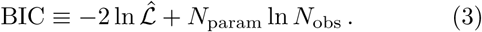

For each criterion, a lower score indicates a better model type. Each criterion effectively quantifies model-type complexity as the number *N* param of independently fitted parameters. The criteria differ in how they weight this complexity (for any reasonable *N* _obs_, BIC weights *N*param more heavily than AIC does) compared to the quality of fit, quantified by the maximum likelihood 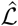 of the model type [33–35] (maximized with respect to all parameters *{θ}*):

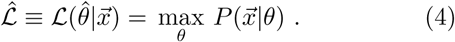

Here 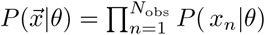 is the probability that the data 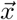 (of dimension *N*_obs_) would be generated by the given model type with parameter values θ, and 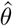 is the likelihood-maximizing parameter set. Modern crystallographic refinement software, such as phenix.refine, use maximum likelihood methods and output a final likelihood estimate along with the conventional metrics (*R*_free_) and the refined structural model.

In this work, we examine the performance of *R*_free_, AIC, and BIC at judging nested pairs of model types that represent common choices crystallographers face during structure refinement. We find that information criteria are more stringent than *R*_free_-based model selection for protein crystallography. On simulated data, AIC and BIC avoid errors that bedevil *R*_free_ at high noise, where *R*_free_ often favors types of models more complex than those used to simulate synthetic data. On experimental data, AIC and BIC rarely prefer more complex model types, exhibiting greater conservatism than *R*_free_. Our results suggest that crystallographers should employ complex model types with greater caution and rely on follow-up biochemical tests when interpreting heterogeneity from these models.

## II. METHODS

We calculated synthetic structure factor amplitudes from a collection of the 50 protein crystal structures studied by Phillips and co-workers using ensemble refinement [14]. We then refined structures against the resulting synthetic and the original experimental data (Fig. 1).

**FIG. 1.**
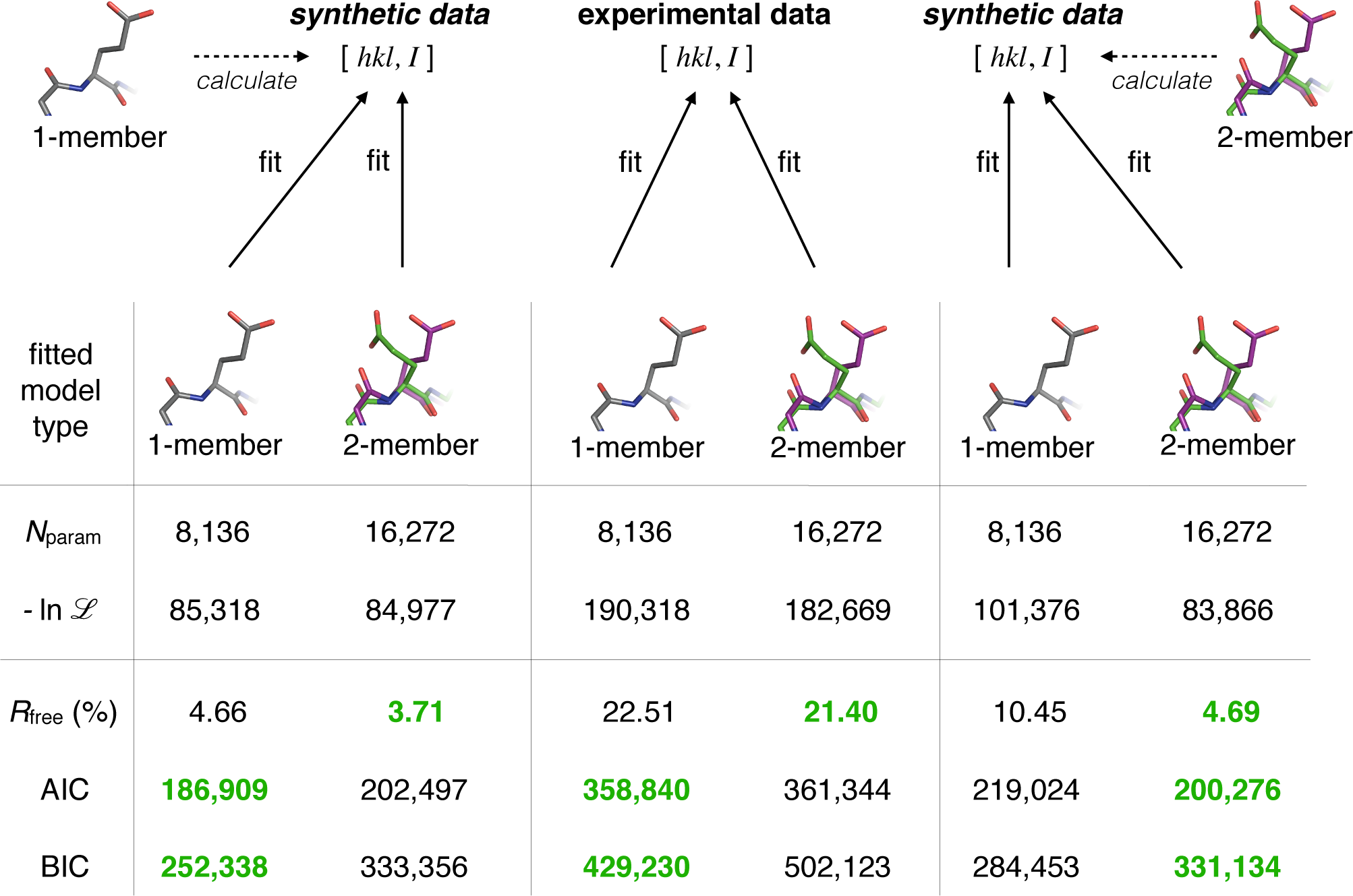
Method schematic. PDB structure (here, a single structure with group isotropic B-factors) is used to generate synthetic structure factor amplitudes, and noise is added (here, 5% of average amplitude). Alternative synthetic data is generated from the complex model type (here, two structures with group isotropic B-factors). The two model types (one or two structures) are fit to each of the synthetic datasets, or to experimental data, generating model selection criteria (*R*_free_, AIC, and BIC) to judge the appropriateness of each model type for the given dataset. Lower criteria scores (corresponding to preferred model type) are in bold green. Numerical results are from 1XY7/2Q48.

We created synthetic data by calculating X-ray amplitudes [36, 37] using phenix.fmodel from models derived from the original PDB entries, but with varying structural heterogeneity, according to several different model types (listed in Table I). Solvent molecules were treated explicitly,*i.e.* with no bulk solvent mask. We then added to each resulting structure factor amplitude a random deviate sampled from a Gaussian distribution, with standard deviation *ξF*_avg_ ranging from a large fraction (ξ = 50%) to negligible fraction (ξ = 0.05%) of the average amplitude *F*_avg_. Resulting negative-valued amplitudes were excluded from further analysis.

The model types varied with respect to three choices faced by crystallographers when deciding the appropriate model-type complexity: whether to fit a distinct individual B-factor for each atom or a single group B-factor for all atoms, whether to fit isotropic or anisotropic B-factors (corresponding respectively to one or six free parameters per B-factor), and whether to fit one or multiple discrete structural conformations. Thus we employed the following model types: one structure with identical isotropic B-factors uniformly fit as a group, one structure with individual isotropic B-factors fit independently for each atom, one structure with individual anisotropic B-factors, and finally fixed-number ensembles of structures with equally weighted occupancies (with number of ensemble members ranging from 1 to *N*_s_, the number in the ensemble PDB based on coordinates deposited by [14]), each with individual isotropic B-factors. For each simulation, we maintained the resolution limits of the original experiment.

**TABLE I.** Number *N*_param_ of parameters for each model type, containing *N*_S_ members and *N*_a_ atoms per member. XYZs: *xyz* coordinates; ADPs: atomic displacement parameters (B-factors).

| $N_s$ | B-factor individuality | B-factor isotropy | XYZs | ADPs | $N_{\text{param}}$ |
| --- | --- | --- | --- | --- | --- |
| 16 | individual | isotropic | $48N_a$ | $16N_a$ | $64N_a$ |
| 8 | individual | isotropic | $24N_a$ | $8N_a$ | $32N_a$ |
| 4 | individual | isotropic | $12N_a$ | $4N_a$ | $16N_a$ |
| 1 | individual | anisotropic | $3N_a$ | $6N_a$ | $9N_a$ |
| 2 | individual | isotropic | $6N_a$ | $2N_a$ | $8N_a$ |
| 1 | individual | isotropic | $3N_a$ | $N_a$ | $4N_a$ |
| 1 | group | isotropic | $3N_a$ | 1 | $3N_a+1$ |

We also made comparisons to the experimental data available from the PDB. For each protein, we downloaded the X-ray structure factors and converted to MTZ file format [36]. As needed, we generated restraint information for geometries of molecules present in PDB models but not already specified by Phenix’s default restraint libraries, using the electronic Ligand Builder and Optimization Workbench (eLBOW) [38]. We neglected anomalous scattering, using only the first array of observed structure factors in each experimental MTZ file.

For each set of synthetic or experimental amplitudes, we performed crystallographic structure refinement using the Phenix platform [26]. We refined the same set of model types as those described above. For each pair of model types, particular constraints on the parameters of the complex model type transform it into the simple one (*i.e.*, the model types are nested). For example, constraining all individual B-factors to be identical produces a group B-factor. Thus, any parametrization (fit) of the simple model type has a corresponding parametrization of the complex model type that generates identical data, hence the best-fit parameters of the complex model type must provide at least as good a fit to data as the best-fit parameters of the nested simple model type.

To maximize the ability of the refinement procedure to find the globally optimal fit (and hence the maximum likelihood as needed for the information criteria), we initialized each refinement for simulated data with the closest possible model to the original model used to create the data (before added noise [39]). This was trivial for fitting the same model as that used to generate the data. To fit an *N*_fit_-member ensemble model type to synthetic data simulated from an *N*_data_-member ensemble model type, when *N*_fit_ < *N*_data_, we initialized the refinement with the first *N*_fit_ members from the ensemble. When *N*_fit_ > *N*_data_, we initialized with *N*_fit_*/N*_data_ copies of each of the *N*_data_ members, with a small random shift added to each coordinate (using the flag “sites.shake=0.01”) to ensure stability of the refinement. Anisotropic refinements of isotropically generated data were seeded with an initial anisotropic model generated using the phenix.pdbtools “convert to anisotropic” command, which changes the B-factor representation from a scalar to a diagonal matrix with diagonal entries all equal to the original single B-factor. Similarly, an isotropic model was generated from an anisotropic model using the Phenix “convert to isotropic” command, which converts the matrix B-factor representation to a scalar that preserves the B-factor averaged over all directions. An initial group B-factor model was generated from an individual B-factor model by replacing each B-factor with the mean value across all B-factors. An individual B-factor model was seeded from a data-generating group B-factor model by giving each atom the same initial B-factor.

We calculated *R*_free_ (1) on 5-10% of the data (omitted from calculating *R*_work_), randomly selected independently of that fraction used for *R*_free_ in [14]. We calculated AIC (2) and BIC (3) using the highest likelihood *ℒ* (defined in Eq. (4) of Ref. [25]) among those Phenix reported during each of 20 macrocycles of semi-automated crystallographic model type refinement [40, 41], using the remaining 90% of working data. Table I lists the number of fitted parameters for each model type. We scored the relative performance for each nested pair of model types according to each criterion *C* ∈ {*R*_free_, AIC, BIC} using

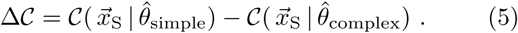

Δ*C* > 0 indicates a preference for the complex model type and Δ*C* < 0 a preference for the simple one.

## III. RESULTS

Synthetic data provide a substantial benchmarking advantage, because the data-generating model and noise level are specified. For a given pair of model types, when the data was generated by a parametrization of the simple model type, that simple model type should always be preferred over the complex one, regardless of the noise level. Thus we compared the performance of nested pairs of model types on synthetic data generated by the simpler one of the pair. Whenever a criterion prefers a model type more complex than that used to generate the data, this indicates an unambiguous model selection error. (Note that the reverse scenario is ambiguous: when data is generated by the complex model type, at zero noise the complex model type should be preferred; however, above some finite noise level—dependent on the true data heterogeneity and generally difficult to determine *a priori*—the simpler model type is the right choice, because the noise prevents faithful reconstruction of extra structural detail by the complex model type.)

We first investigated a modest increase in complexity between grouped and individual isotropic B-factors. This would increase the number of parameters from 1,801 to 2,400 on a 600-atom (~100-residue) protein. Conventionally, individual B-factors are refined at better than ~3.5*Å* resolution. Figure 2 shows the difference Δ*C* ≡ *C*_group_ − *C* individual, and thus the preference of different criteria for group or individual B-factor model types on data simulated from either the simpler (group B-factor) model type or the more complex (individual B-factors) model type. As noise increases, all methods tend to favor the simple model type. As data heterogeneity increases from group to individual B-factors, all methods look more favorably on the complex model type. As resolution decreases, all methods show a weak trend toward an increasing preference for the simple model type. Of most significance, AIC and BIC never choose the wrong model type (one more complex than the one generating the data), but at high noise *R*_free_ sometimes does.

**FIG. 2.**
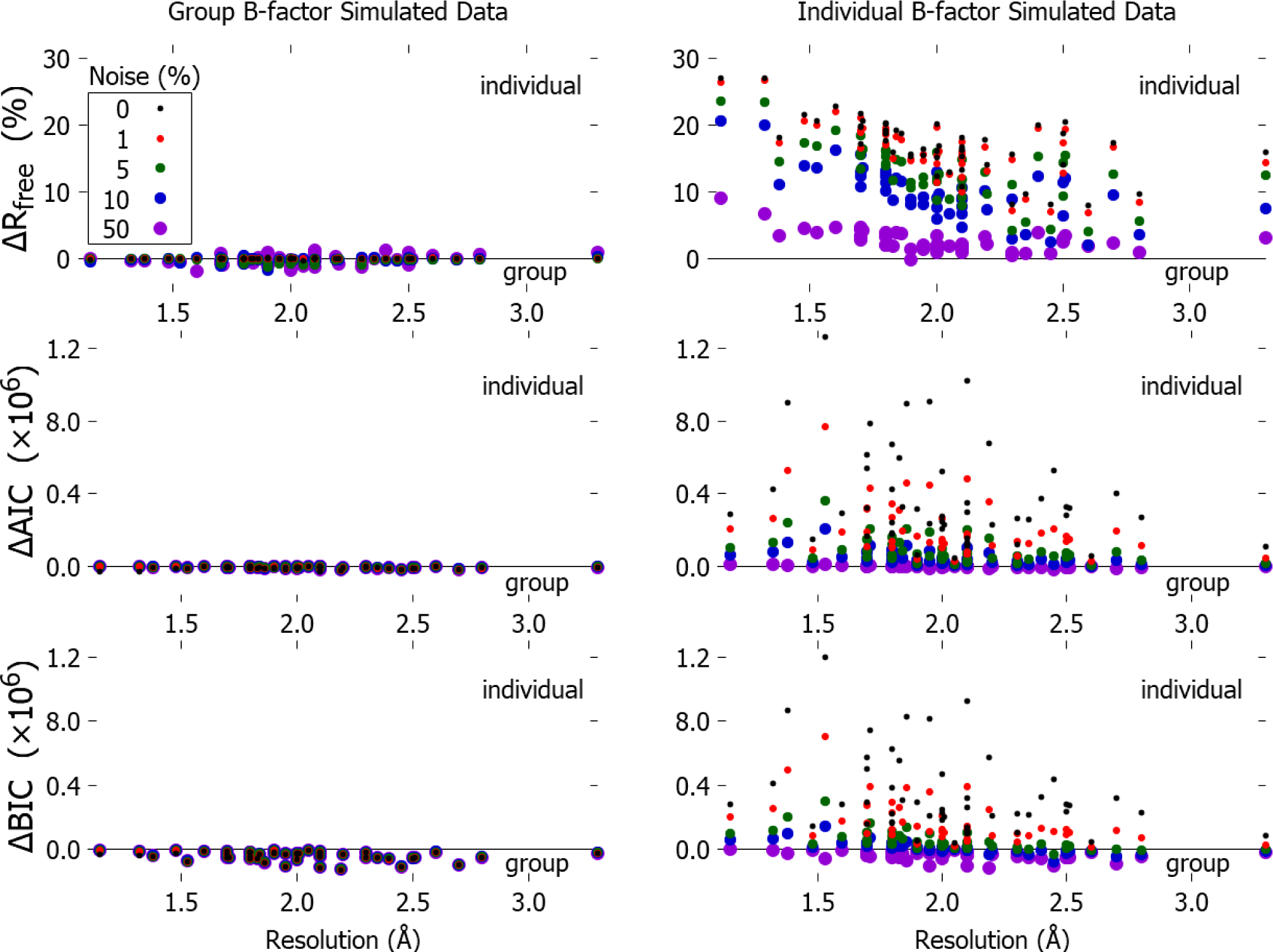
*R*_free_, unlike AIC and BIC, is vulnerable to erroneously preferring an individually fit B-factor model type. Preference Δ*C* ≡ *C*_group_ − *C*_individual_ for complex model type (individual fit B-factors) over simple model type (group fit B-factor) as a function of resolution, for *C* = *R*_free_, AIC, and BIC (top row to bottom row). Higher value indicates higher preference for complex (individual) model type, with zero indicating indifference. Left column: data generated from model type with single group B-factor. Right: data generated from model type with individual B-factors. Noise varies from none (small black points) to high (standard deviation 50% of average amplitude, large purple points).

Next, we examined the switch from isotropic to anisotropic refinement, which is conventionally performed only at high resolution (*i.e.*, better than ~1.2-1.5 *Å* resolution) [42]. Figure 3 shows the preference of different criteria for isotropic or anisotropic B-factor model types on data simulated from either the simpler (isotropic) model type or the more complex (anisotropic) one. As noise increases, AIC and BIC increasingly prefer simpler model types, but *R*_free_ sometimes prefers more complex ones. As data heterogeneity increases from isotropic to anisotropic B-factors, all methods show greater preference for the complex model type. As resolution decreases, all methods increasingly prefer simpler model types. AIC and BIC never choose the unambiguously wrong model type, but at medium noise *R*_free_ often does.

**FIG. 3.**
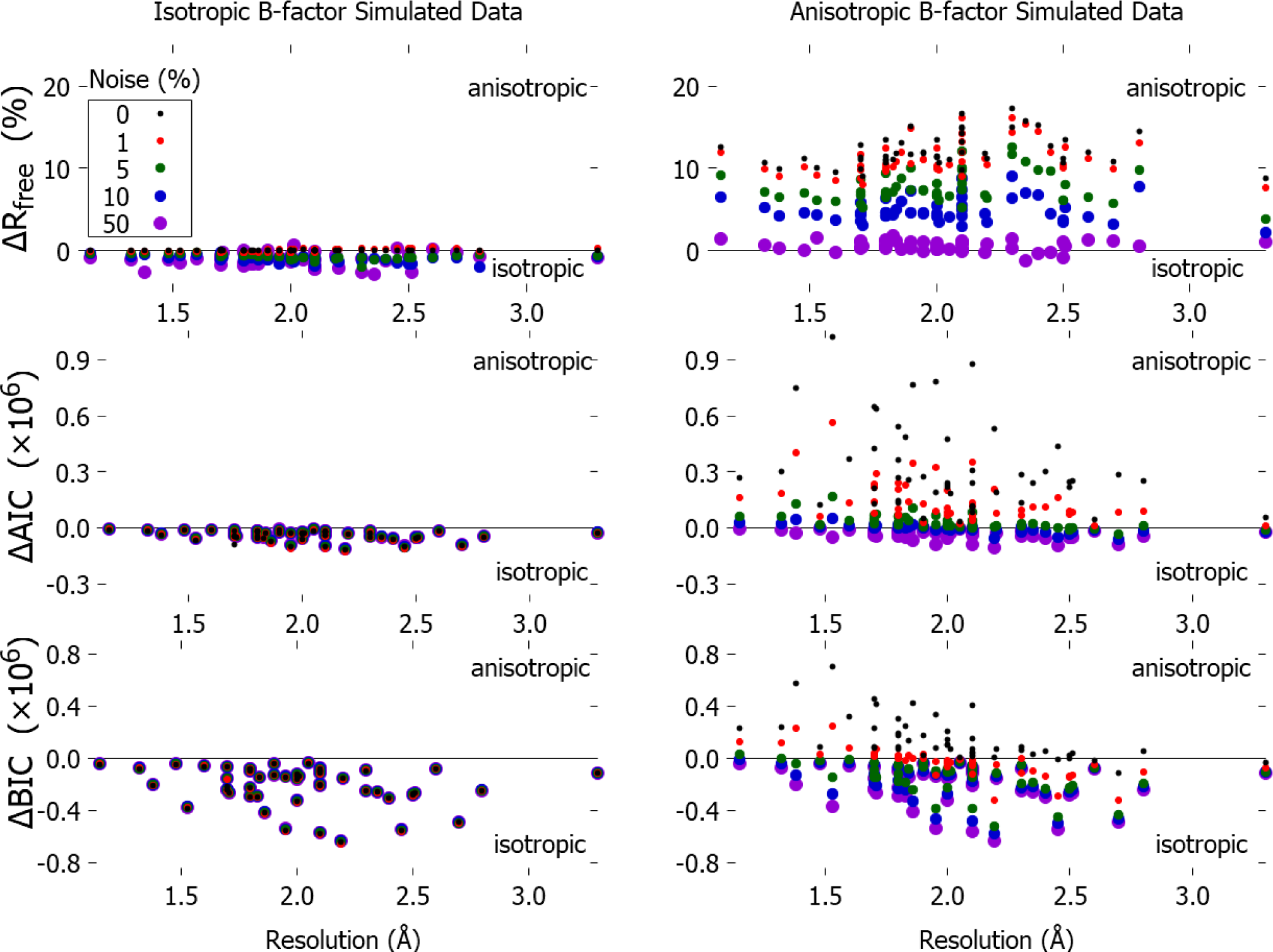
Preference Δ*C* ≡ *C*_iso_ − *C* aniso for complex model type (anisotropic B-factors) over simple model type (isotropic B-factors) as a function of resolution, for *C* = *R*_free_, AIC, and BIC (top row to bottom row). Higher value indicates higher preference for complex (anisotropic) model type, with zero indicating indifference. Left column: data generated from isotropic model type. Right column: data generated from anisotropic model type. Noise as in Fig. 2.

Although anisotropic B-factors can fit elliptical electron density distributions, multiple conformations are needed to fit distributions with multiple minima. For this reason, various ensemble refinement methods have been developed. Here we used fixed-number ensembles similar to those created by Phillips and colleagues in [14]. Figure 4 shows the preference of different criteria for model types containing fewer or greater numbers of ensemble members, on data simulated from the simpler (fewer ensemble member) model type. As noise increases, *R*_free_ sometimes prefers more complex model types, AIC rarely does, and BIC never does. As resolution decreases, *R*_free_ shows no strong trend, but AIC and BIC increasingly prefer simpler model types. BIC never erroneously chooses the overly complex model type, but AIC on rare occasions does, and even at low noise *R*_free_ frequently does.

**FIG. 4.**
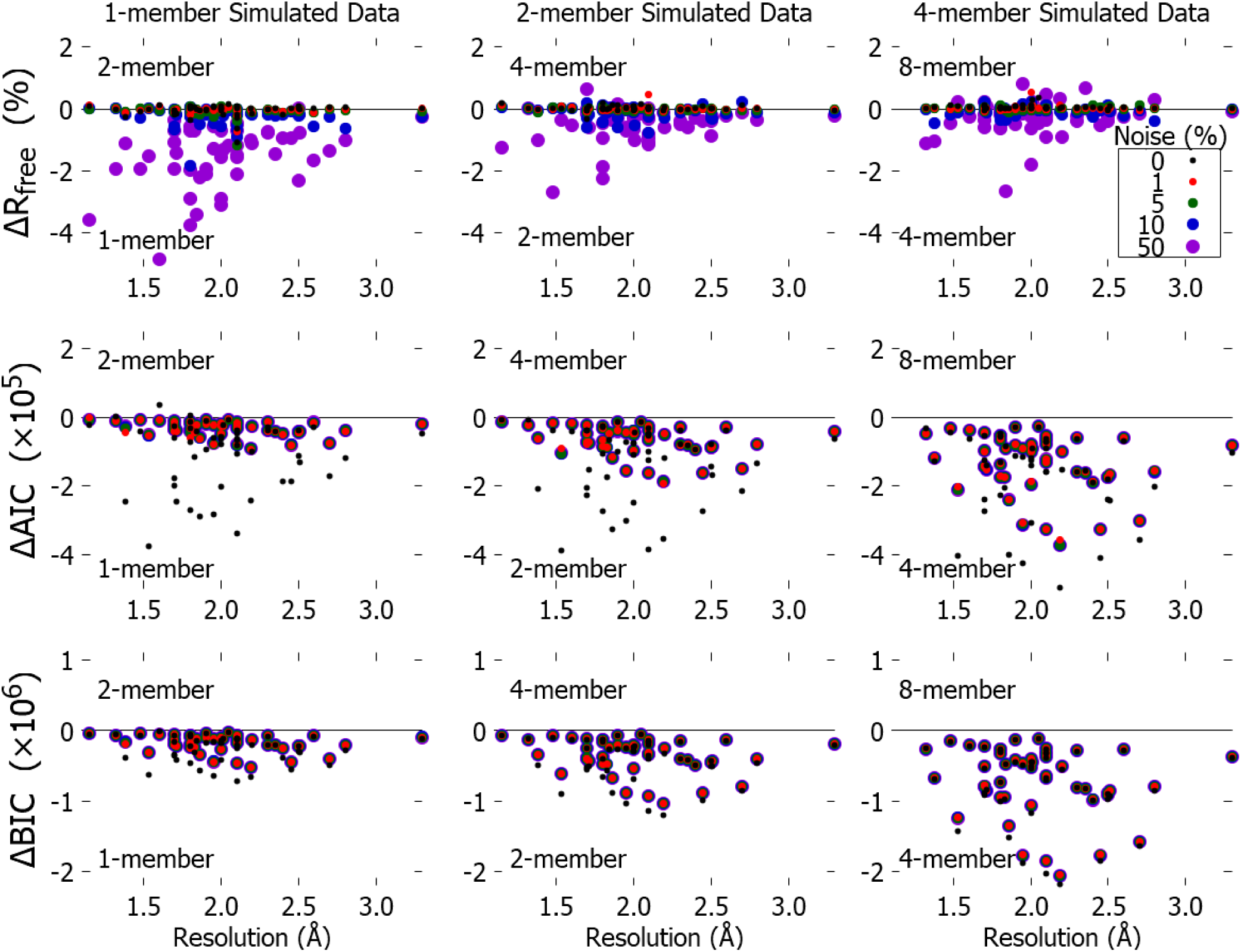
Preference Δ*C* ≡ *C*_*N*_ − *C*_2*N*_ for complex model type (twice as many ensemble members as used to generate data) over simple model type (same number of members) as a function of resolution, for *C* = *R*_free_, AIC, and BIC (top row to bottom row). Higher value indicates higher preference for complex model type (with more members), with zero indicating indifference. Data generated from model type with one (left column), two (middle), or four members (right). Noise as in Fig. 2.

Figure 5 summarizes, as a function of noise, the proportion of ‘false positives,’ where a selection criterion prefers the complex model type when the synthetic data is generated from the nested simple model type. (It is worth noting that typical noise estimates for successful diffraction experiments are likely to be in the 1-10% range [43].) In general, across the three nested comparisons shown here, AIC and BIC make more conservative selections, consistently identifying simple data sets and rejecting the overfit model types that sometimes fool *R*_free_. To explicitly address the possibility of *R*_free_ overfitting, we introduced an additional selection criterion, the heuristic ‘supplemented *R*-factor’*R*_supp_ that only prefers the complex model type if *R*_free_ is lower *and* the *R*_free_–*R*_work_ gap hasn’t dramatically increased ((*R*_work_ − *R*_free_)_complex_ < 1.5(*R*_work_ − *R*_free_)simple).*R*_supp_ achieves the same low false-positive rates as AIC and BIC, except in the isoaniso comparison at low-to-moderate noise.

**FIG. 5.**
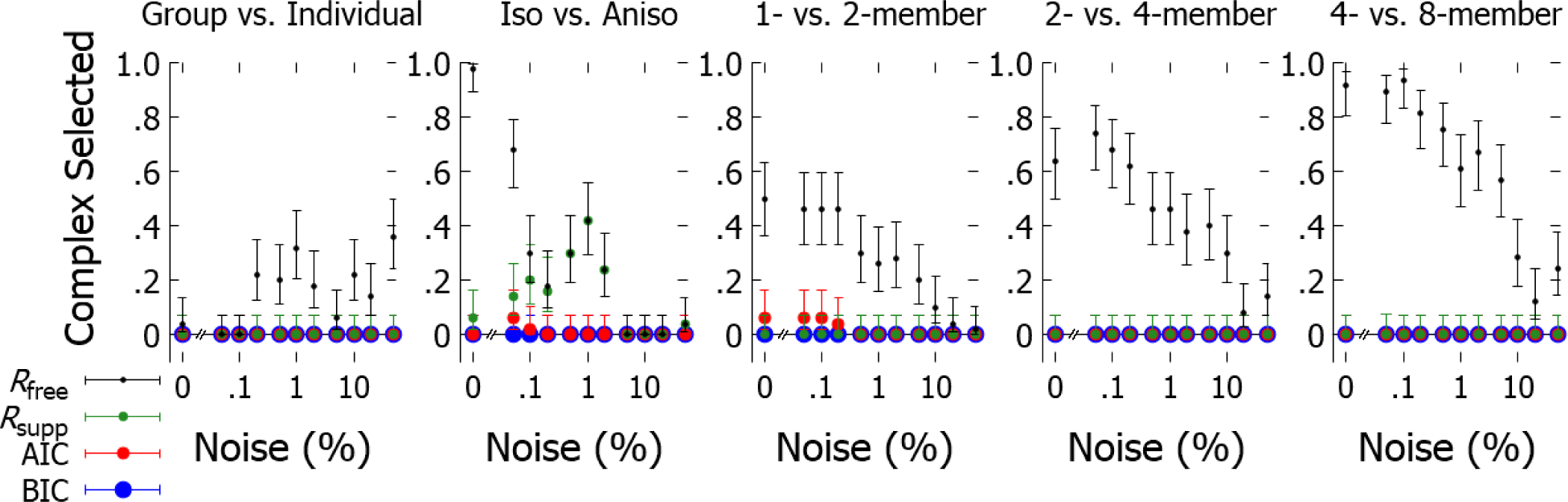
Proportion of proteins where selection criterion (*R*_free_,*R*_supp_, AIC, or BIC) prefers a complex model type to fit data generated from a simple model type, as a function of noise. Simple and complex model types compared are listed above each panel. Error bars show Wilson score 95% confidence interval.

Next, we examined the experimental data sets from Levin,*et al.* [14] where the ‘true’ model type is not known, and many other errors (*e.g.*, measurement errors or initial model errors) may be conflated. Single-member (simple) and eight-member (complex) ensembles were previously deposited in the PDB, which allowed us to evaluate *R*_free_, AIC, and BIC for model selection. In addition, we calculated simulated data based on the single-member or eight-member ensembles. Figure 6 (left column) shows that *R*_free_ evaluates many of the complex ensembles as better fits to the experimental data. However, both AIC and BIC consistently prefer the simpler model type. This result against experimental data is corroborated by the results against data simulated from a single-member ensemble (middle) and from an eight-member ensemble (right). Note that the experimental results are comparable to simulated results for the synthetic noise levels used here, lending some credibility to the incorporation of Gaussian noise in the calculated data. From single-member simulated data, AIC and BIC always prefer the simpler model type. Similarly, the information criteria prefer the simpler model type even when the underlying data are calculated from the complex model type, except for AIC in some cases with very low noise.

**FIG. 6.**
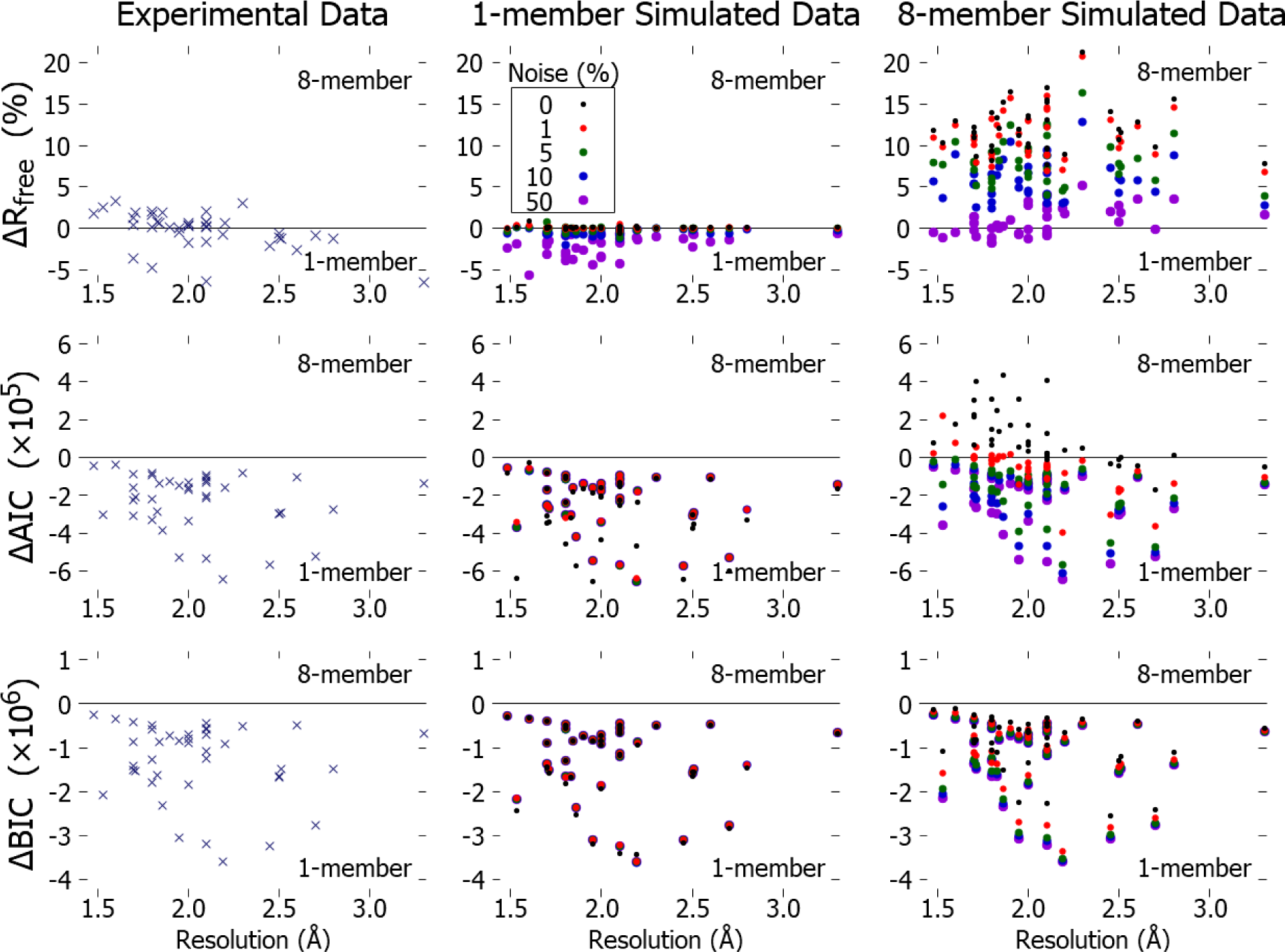
Model selection on synthetic data makes similar judgments to model selection on experimental datasets. Preference Δ*C* ≡ *C*_1−member_ − *C*_8−member_ for complex (8-member) model type over simple (1-member) model type as a function of resolution, for *C* = *R*_free_, AIC, and BIC (top row to bottom row). Higher value indicates higher preference for complex model type (with more members), with zero indicating indifference. 1- and 8-member ensemble model types are fit to experimental diffraction patterns (left column,*×*), or to synthetic patterns (filled dots, *•*) simulated using one member (middle column) or eight members (right column). Noise as in Fig. 2.

**FIG. 7.**
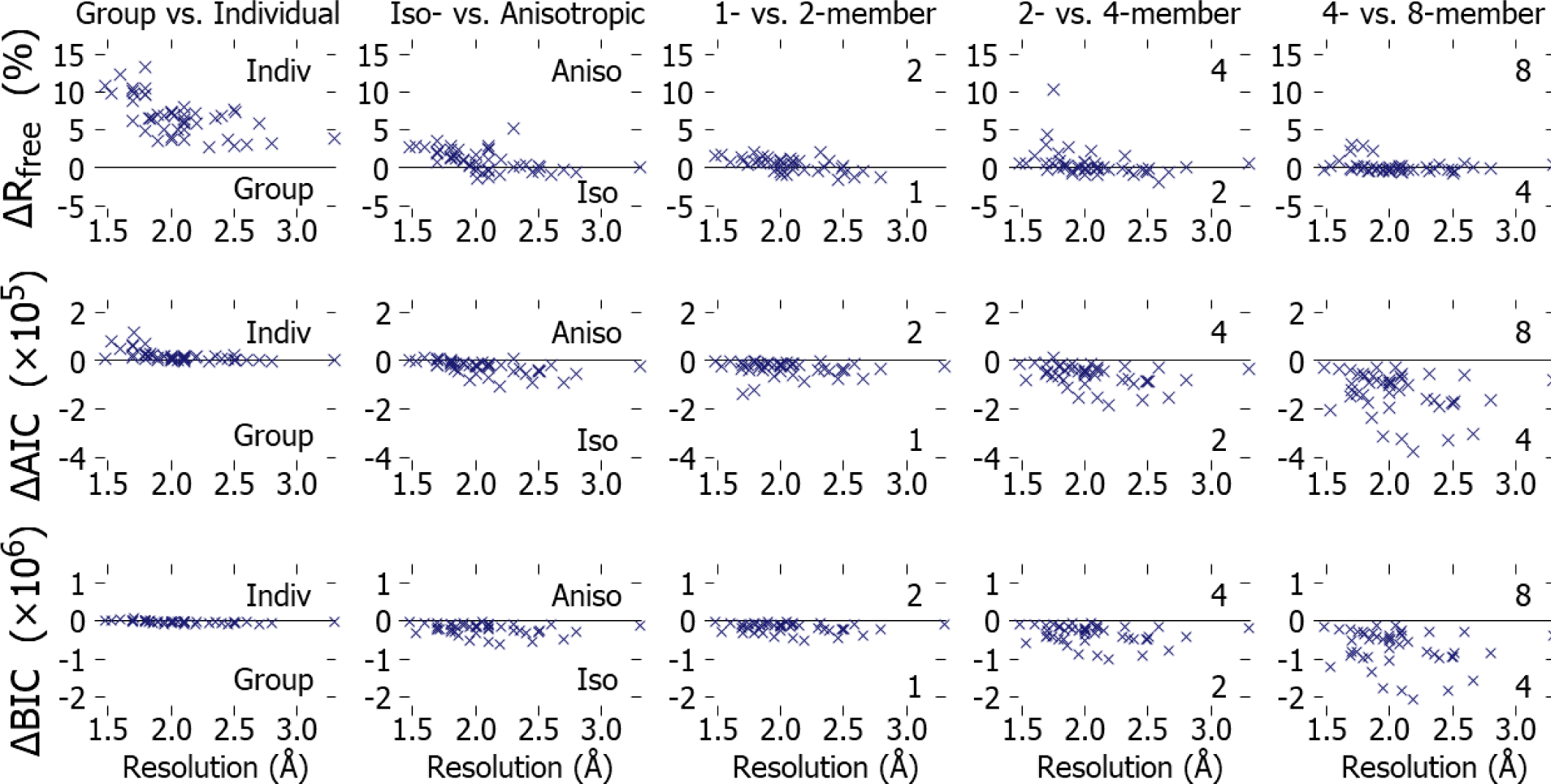
Preference *C* ≡ *C*_simple_ − *C*_complex_ for complex model type over simple model type to fit experimental data, as a function of resolution, for *C* = *R*_free_, AIC, and BIC (top row to bottom row). Higher value indicates higher preference for complex model type, with zero indicating indifference. Columns (left to right) represent model-type comparisons of group vs. individual, iso- vs. anisotropic, 1- vs. 2-member, 2- vs. 4-member, and 4- vs. 8-member, respectively. Noise as in Fig. 2.

Next, we performed refinements of different nested model types against the experimental data of Levin,*et al.* and evaluated how often the more complex model type was selected (Fig. 8). For group vs. individual B-factors, only BIC rejects the more complex model types. This suggests that BIC is more stringent than common practices in the field, where individual B-factors are normally refined across the entire range of resolutions used here. Interestingly, although anisotropic B-factors were preferred for the majority of models by *R*_free_, AIC only preferred them for a small number of models. To judge whether this is consistent with heuristics commonly employed in the field, we assessed whether the *R*_free_-*R*_work_ divergence had dramatically increased for some of the models refined with anisotropic B-factors. Indeed, Fig. 8 also shows there was evidence of overfitting for many of these model types as detected by the *R*_supp_ criterion. In contrast, for ensemble model types,*R*_free_ and *R*_supp_ tracked closely, and AIC rarely (and BIC never) preferred the more complex model types.

**FIG. 8.**
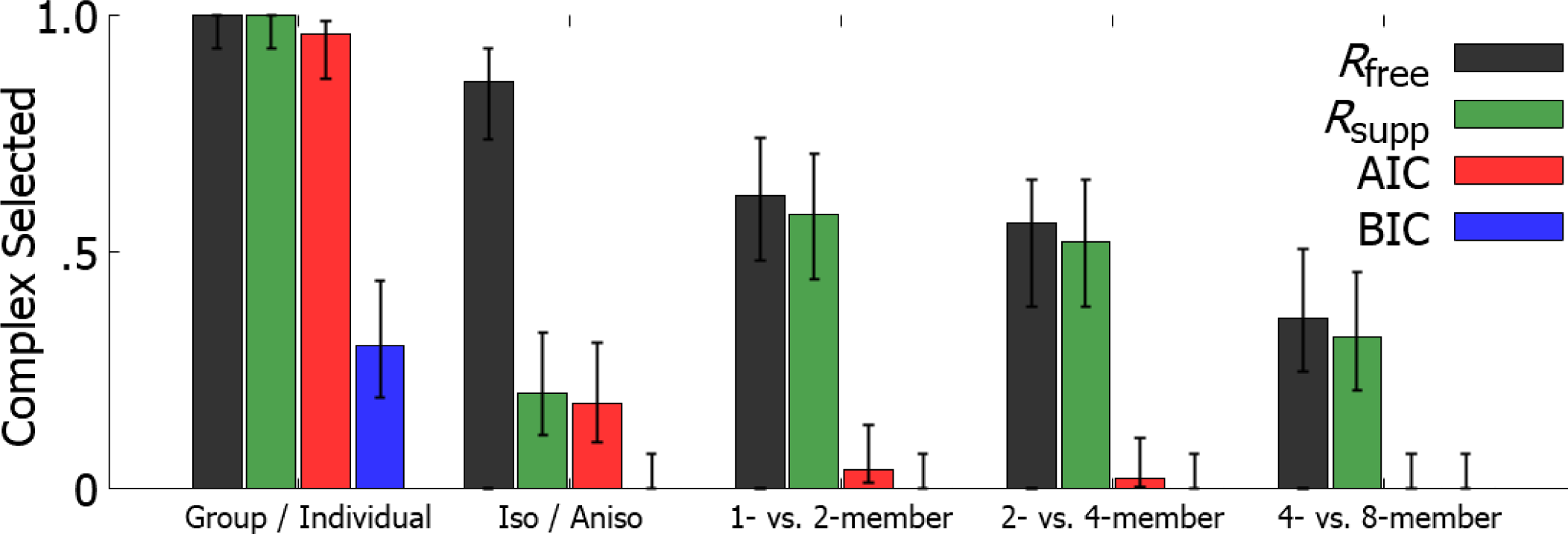
Proportion of proteins where selection criterion *C* (*R*_free_, *R*_supp_, AIC, or BIC) prefers a complex model type to fit experimental data. Error bars show Wilson score 95% confidence interval.

## IV. DISCUSSION

Protein crystallography has historically developed model types with increasingly detailed representations of conformational heterogeneity in protein crystals [14, 44–48]. Yet it remains unclear how to know when heterogeneous model types are justified by the experimental data. A complex model type with many parameters may provide a better fit to data than a simple model type with few. Yet, parameters added to improve an existing model type’s quality of fit bring with them the hazard that they may not parsimoniously describe the physical characteristics of interest. The practical trade-off between accuracy (quality of fit) and parsimony (reduced vulnerability to overfitting) defines the long-standing problem of model selection: What is the most appropriate amount of complexity to use to model a particular experimental dataset?

*R*_free_ is often incorporated into model selection methods [49, 50]. But if *R*_free_ is exploited both to select model type and to estimate model type errors [51], it may fall prey to precisely the problem it was designed to address: selecting an overly complex model type to fit spurious characteristics of ambient noise. Adding consideration of the *R*_work_-*R*_free_ gap (in this work *via R*_supp_) can reduce this overfitting, warranting further study of its role in model selection, but current uncertainty on how best to incorporate both *R*_work_ and *R*_free_ presents practical complications.

As an alternative for model selection, information criteria mitigate the risk of overfitting superfluous model type parameters by assigning explicit penalties to discourage use of complex model types. Information criteria thus explicitly weigh the risks of additional parameters against the benefits of using them.

We find that AIC and BIC are credible criteria for model selection in protein X-ray crystallography, outperforming *R*_free_ on data simulated from simple model types and suggesting caution when employing more complex model types in refinement against experimental data. In synthetic data generated from the simpler model type— where one should never prefer the more complex model type, regardless of noise level—*R*_free_ often erroneously prefers the unambiguously wrong, more complex (overfit) model type. As synthetic noise increases, information criteria more predictably (compared to *R*_free_) increase their preference for simpler model types (Figs. 2,3,4). On experimental scattering data where we do not *a priori* know the correct answer, AIC and especially BIC judge model types significantly more conservatively than does *R*_free_ (Fig. 6). Whereas *R*_free_ on average supports the use of more complex model types—across all our pairwise comparisons—to fit the experimental diffraction data considered, AIC and BIC generally prefer simpler model types (Fig. 8).

As a measure of quality of fit, we used Phenix’s likelihood function. Our work assumes that this likelihood quantifies the probability of generating the observed data from the proposed parameterization of the model type and noise model.

Knowledge about typical protein geometry is incorporated into structure refinement in the form of what is effectively a Bayesian prior over bond lengths, bond angles, torsion angles, clashes, B-factor variation among neighboring atoms, and so on [52]. These various restraints on geometry and B-factor variation reduce the effective number of free parameters during refinement [31], but are not accounted for in our parameter counts (Table I). As complex model types have more parameters and hence will have their effective number of free parameters reduced more than simple model types, the information criteria (as currently formulated) may be overestimating the complexity of the complex model types, and thus overly penalizing them. Additional complications may emerge because the weight between restraints and data is often optimized based on *R*_free_.

Moreover, refinement should be less likely to find the best-fit parameters for a more complex model type, given the higher-dimension parameter space to search through. Thus standard refinement can be expected to underestimate the quality of fit possible for a complex model type.

Our noise model is highly simplified and thus results in *R*-factors that refine to convergence. Such small *R*_free_values are not observed for real data, where the ‘*R-*factor gap’ can be sizable [53]. A more sophisticated noise model, such as that incorporated in MLFSOM [53], could alleviate this discrepancy.

The multiconformer model type [54], which has alternative conformations for some but not all atoms or residues, represents a middle ground between single-member and multi-member ensemble model types. Future study using our model selection framework may illuminate its potential advantages. However, the multi-conformer model type presents additional practical difficulties surrounding the choice of how many atoms—and which ones—merit representation by alternative conformations, and ambiguity about the true number of free parameters that are fitted since it considers potentially all atoms or residues for more complex representation. Torsion-angle refinement [55] is another method that reduces the number of free parameters while attempting to retain sufficient flexibility to capture true structural heterogeneity.

This work represents an initial investigation of existing model selection techniques, directly applied to judging protein model types in fits to X-ray crystallography data. For the most part, we have treated Phenix as a black box; taking the likelihood it reports at face value, as an end user would. An interesting extension would relax the simplifying assumption embodied in AIC and BIC that the posterior distribution is unimodal (with a single peak in probability), allowing for the multimodal nature of the prior protein knowledge (*e.g.*, the Ramachandran plot has multiple peaks), which becomes especially relevant at the relatively low parameter-to-observation ratios here. One could also sample over the posterior distribution (such as in the Deviance Information Criterion) [56] instead of making the strong assumptions AIC and BIC do about the shape of the posterior distribution.

This framework is quite general and should also provide an interesting perspective on model selection in other structural biology contexts such as nuclear magnetic resonance, cryo-electron microscopy, and integrative modeling [57]. Indeed, X-ray crystallography likely represents a best-case scenario in terms of the ratio of data to free parameters and relatively low noise, so problems encountered in overfitting crystallographic models to data should only be more prominent in more data-sparse structural fields.

## Appendix A: Likelihood

Here we detail Phenix’s likelihood [25]. The likelihood assumes that the joint probability distribution *P*(*{F*_s_}) describing the structure factor amplitudes {*F*_s_} indexed by reciprocal vectors s in the set *S* = {s} may be approximated by a product of independent distributions,

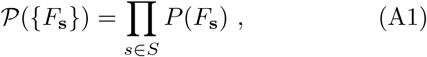

with the probability of a model producing an individual amplitude assumed to be Gaussian in the modulus of the discrepancy between prediction and data,

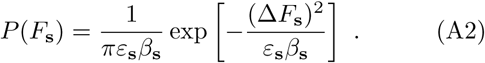

Here 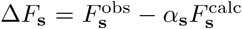 quantifies the difference between the observed amplitude 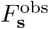 and the predicted amplitude 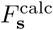.

The fitted parameters α_s_ and β_s_ quantify the expectation and variance of the amplitude error [58]. The fitted multipliers ε_s_ adjust for the varying mean intensities of the different reflections [24]. Parameters α_s_, β_s_, and *ε*_s_ are all fitted during the refinement, but their number do not vary among the physical model types. So when comparing model type performance, subtracting one information criterion from another, these contributions to the number of fitted parameters cancel.

For acentric reflections [25], integrating Eq. (A2) over the (inaccessible) complex phase information [25] gives the distribution of amplitudes

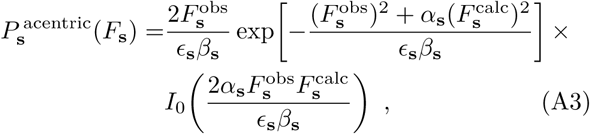

for *I*_0_(*x*) the modified Bessel function of the first kind. For our purposes, it is sufficient to neglect the model-independent terms in Eq. (A3) to obtain the model-dependent (negative) log-likelihood for each reflection

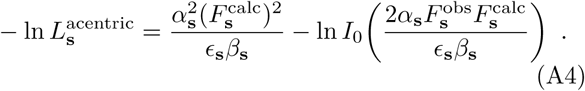

For centric reflections [25], Eq. (A2) instead reduces to

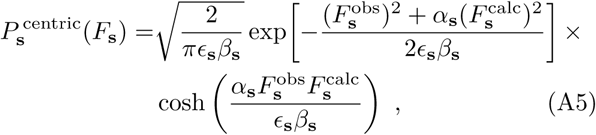

giving the model-dependent terms in the (negative) log-likelihood *per* reflection,

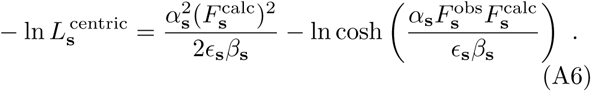

## ACKNOWLEDGMENTS

The authors thank Dave Campbell and Biljana Jonoska-Stojkova (SFU Statistics), Aidan I. Brown (SFU Physics), James Holton (Lawrence Berkeley National Lab Biosciences), Tom Terwilliger (Los Alamos National Lab), George Phillips and Mitchell Miller (Rice Biosciences) for illuminating conversations and feedback on the manuscript, and especially Pavel Afonine (Lawrence Berkeley National Lab) for his continuing support with the Phenix refinement engine. This research is supported by NIH Grant GM110580 (J.S.F and D.A.S.), a Tier-II Canada Research Chair (D.A.S.), and WestGrid (www.westgrid.ca) and Compute Canada Calcul Canada (www.computecanada.ca).

